# Isoform-specific nicotinic responses are present in distinct subtypes of prefrontal layer V pyramidal neurons

**DOI:** 10.1101/2023.04.10.536291

**Authors:** Ashutosh V. Patel, Anthony Nguyen, Pietro Paletta, Elena Choleris, Craig D.C. Bailey

## Abstract

The neurotransmitter acetylcholine supports goal-directed cognitive functions via activation of its nicotinic and muscarinic classes of receptors within the prefrontal cortex. These receptors are expressed on pyramidal neurons located within layer V of the prefrontal cortex, which integrate afferent signals and contribute toward cognitive circuits via efferent projections to cortical and subcortical targets. Using whole-cell electrophysiology, retrograde labelling, and neuron reconstruction in the juvenile mouse prefrontal cortex, we identified three unique isoform-specific nicotinic receptor responses that are present in distinct subtypes of layer V pyramidal neurons. Broadly, we observed α7 or α7/β2* nicotinic responses in burst-firing neurons that project to the contralateral cortex or nucleus accumbens, respectively, and β2* nicotinic responses in regular-firing neurons that project to the ventromedial thalamus. These findings provide insight into a receptor isoform-specific mechanism by which nicotinic acetylcholine neurotransmission may support cognitive functions via modulation of distinct efferent projections from this brain region.

## Introduction

The modulatory neurotransmitter acetylcholine (ACh) acts within the rodent medial prefrontal cortex (mPFC) to support goal-directed cognitive functions such as working memory and attention^1–3^. The manner by which ACh fulfills this role is sophisticated and likely multimodal^4,5^, as evidence exists for both a sustained tonic increase in extracellular ACh content^6,7^ and a transient phasic increase in synaptic ACh content^1,2^ for animals performing visual attention tasks. ACh activation of its nicotinic class of receptor (nAChR) plays an important role in this process. For example, the pharmacological or genetic disruption of nAChR function impairs performance^3,8–12^ and the pharmacological augmentation of nAChR function enhances performance^12–14^ for mPFC-dependent tests of cognitive function. The nAChR comprises a group of pentameric ligand-gated ion channels that are permeable to monovalent and divalent cations. The two predominant nAChR isoforms within the mPFC are homomeric α7 nAChRs (composed of five α7 subunits) and heteromeric β2* nAChRs (composed of α4 and β2 subunits with the potential incorporation of one α5 subunit). These isoforms possess distinct functional properties and are differentially expressed within cellular layers and neuron types of the mPFC (reviewed in Abbondanza and colleagues^15^). Within layer V of the mPFC, it has been observed that pyramidal neurons express α7 nAChRs and that both fast-spiking and non-fast spiking interneurons may express combinations of α7 and/or β2* nAChRs^16,17^. However, evidence suggests that layer V pyramidal neurons within other cortical regions express both α7 and β2* nAChRs^18^. It is important to determine whether layer V pyramidal neurons within the mPFC also express both receptor isoforms because this will further our understanding of the sophisticated manner by which cholinergic neurotransmission supports mPFC-based cognitive circuits and behaviours.

Layer V pyramidal neurons integrate inputs from cortical and subcortical sources, placing them in an important position to contribute toward cognitive circuits. They are a heterogeneous population based on electrophysiological and morphological properties. Previous research over the past four decades has characterized layer V pyramidal neurons in multiple cortical regions, allowing for the discovery of distinct neuron subtypes having specific action potential firing patterns, apical dendrite morphologies, and axonal projection targets. In response to depolarizing current injection, action potential firing patterns are described as intrinsically burst-firing (BF) or regular-firing (RF), with sub-variations being reported within each of these categories^19–34^. Although precise terminology can vary depending on the study, BF neurons commonly are described as exhibiting true all-or-nothing bursts of action potentials^19–21,26,33^, or two-to-four action potentials with a markedly short interspike interval (ISI) at the beginning of a train of action potentials^22–25,29,30,32^. RF neurons exhibit a relatively longer, sometimes adapting ISI throughout a train of action potentials^19–30,32^. Other properties of BF and RF neurons are often found to distinguish them into groups having distinct apical dendrite morphologies and axonal projections. In sensory and motor cortices, BF neurons typically have “wide” or “thick” apical dendrite tufts and are “pyramidal tract” (PT) neurons that project to subcortical regions including the striatum, thalamus, spinal cord, pons, and superior colliculus whereas RF neurons typically have “narrow” or “thin” apical dendrite tufts and are “intratelencephalic” (IT) neurons that project to the contralateral cortex and striatum^20,23,25,30,33,35–41^. These properties do not align perfectly in all reports, suggesting potential variation based on sex, age, or recording conditions^23,24,37,42,43^. Moreover, properties of these broad neuron subtypes may differ between sensorimotor and associative cortical regions^34^. In the mPFC, combinations of BF trains, IT/striatal projections, and narrow/thin apical dendrite tufts are typically observed while combinations of RF trains, PT projections, and wide/thick apical dendrite tufts are typically observed^32,34,44,45^, although this is not always found for all properties^31,32^ and both BF and RF neurons have been identified in PT and IT populations in unspecified frontal cortex^27,28^

In this present study, we identify and characterize unique isoform-specific nicotinic responses in three distinct pyramidal neuron subtypes within mPFC layer V sampled from juvenile mice of each sex. Like previous work in mPFC described above, each of these neuron subtypes has its own combination of action potential firing pattern, apical dendrite morphology, and projection target(s). Specifically, we show that α7 nAChR responses are present in BF neurons that have narrow (male) or wide (female) apical dendrite tufts and project to the contralateral mPFC, a combination of α7 and β2* nAChR responses are present in BF neurons that have wide (male) or narrow (female) apical dendrite tufts and project to the ipsilateral ventral striatum (nucleus accumbens), and β2* nAChR responses are present in RF neurons that have wide (male) or narrow (female) apical dendrite tufts and project to the ipsilateral ventromedial thalamus. This work adds the dimension of nicotinic response type and sex differences to the known properties of mPFC layer V pyramidal neuron subtypes, and provides insight into an nAChR isoform-specific mechanism by which cholinergic neurotransmission may influence efferent signalling from this brain region.

## Results

### Three unique nicotinic responses are present in mPFC layer V pyramidal neuron subtypes

Pyramidal neurons within layer V of the mPFC may be categorized into two broad subtypes based on their action potential firing patterns in response to depolarizing current injection. Under our recording conditions in juvenile mice, the injection of 200 pA depolarizing current from rest for 500 ms identified a subtype of BF neurons based on a short initial ISI between the first two action potentials of <10 ms and a subtype of RF neurons based on a relatively longer ISI between the first two action potentials of >10 ms (Fig. 1). The BF subtype may be further divided based on the nicotinic response types that were observed. Pressure application of 1 mM ACh for 100 ms elicited one of two nicotinic responses in BF neurons, which had either a fast rise and fast decay (Fig. 1a_2_) or a fast rise and two-phase fast plus slow decay (Fig. 1b_2_). Responses with a fast rise and fast decay were not affected by the β2* nAChR antagonist dihydro-β-erythroidine (DHβE) but were inhibited completely by the α7 nAChR antagonist methyllycaconitine (MLA) (Fig. 1a), which is consistent with previous reports that α7 nAChRs are present on cortical layer V pyramidal neurons^16,18^. Responses with a fast rise and two-phase decay had fast components inhibited by MLA and the remaining slow components inhibited by DHβE (Fig. 1b), suggesting the presence of both α7 and β2* nAChRs, and possibly also the presence of α7β2 heteromeric nAChRs containing both α7 and β2 subunits (reviewed by Wu and colleagues^46^; referred to herein as α7/β2* responses). We did not test for the presence of true α7β2 heteromeric nAChRs in these neurons because the subunit-selective antagonists MLA and DHβE also inhibit this receptor isoform^47,48^. This same ACh application to RF neurons consistently elicited responses with a slow rise and slow decay that were characteristic of β2* nAChRs^3,49^. This was confirmed pharmacologically, as these responses were unaffected by MLA but were inhibited completely by DHβE (Fig. 1c). Contingency analysis revealed that approximately half of recorded neurons were BF and modulated by α7 responses only (male: 29/57; female: 35/63), approximately one-quarter of neurons are BF and modulated by α7/β2* responses (male: 13/57; female: 15/63), and approximately one-quarter of neurons are RF and modulated by β2* responses only (male: 15/57; female: 13/63) (Fig. 1d). These distributions did not differ between sexes (Chi-square test, *p* = 0.76).

**Fig. 1.**
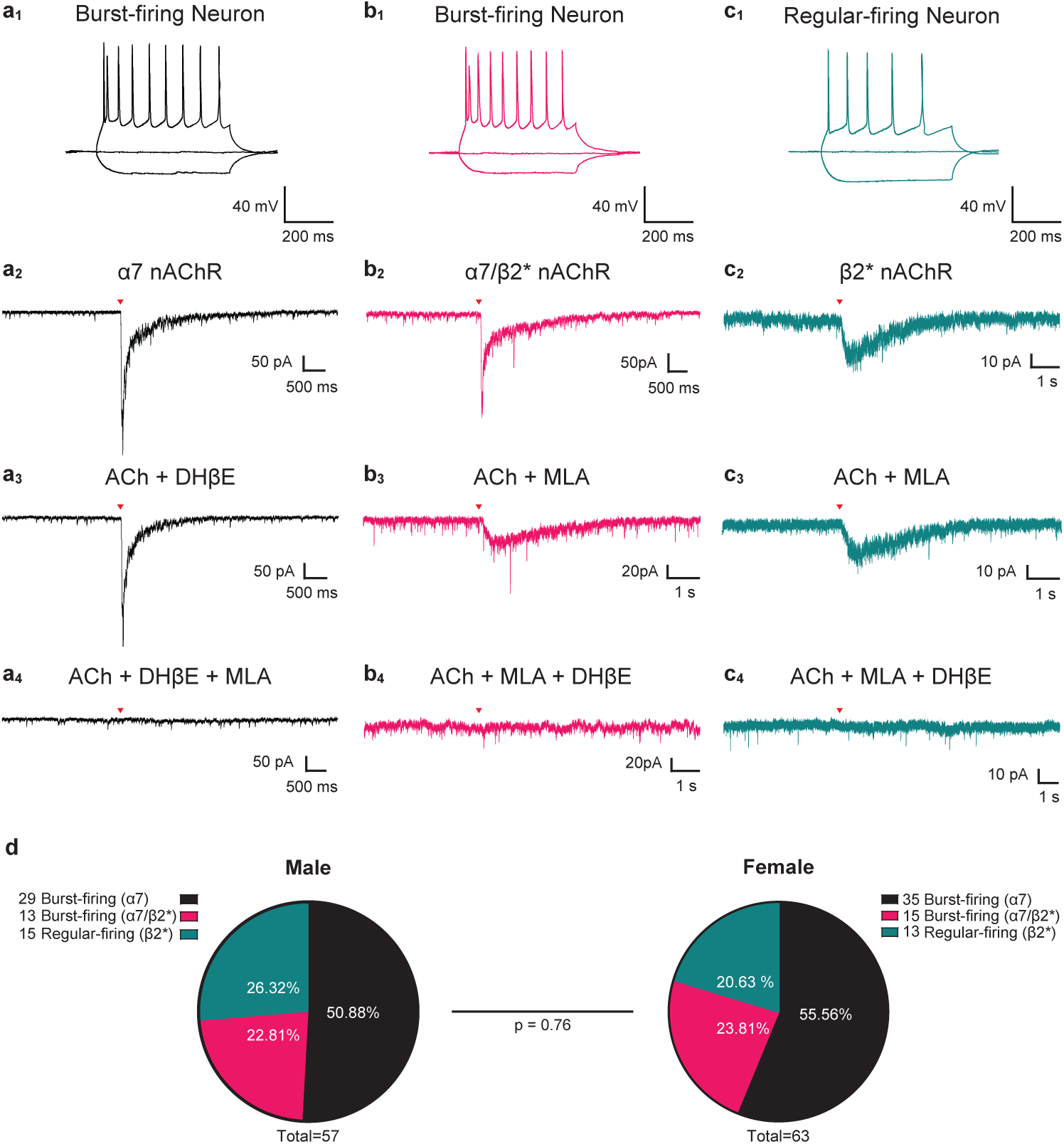
Nicotinic responses observed in prefrontal layer V pyramidal neurons. Pyramidal neurons were sampled randomly from layer V of the juvenile mouse mPFC and recorded for intrinsic electrophysiological properties and nicotinic acetylcholine responses. Three unique nicotinic responses were observed within distinct neuron subtypes. **a** The most common nicotinic response was found in neurons that exhibited an initial burst-firing action potential firing pattern in response to the injection of 200 pA positive current (**a_1_**). This nicotinic response to 100 ms application of 1 mM ACh (indicated by the red arrowhead in **a_2_**) had fast rise and decay kinetics characteristic of homomeric α7 nAChRs, which was confirmed by the lack of effect of 3 μM DHβE (**a_3_**) and the complete inhibition by 10 nM MLA (**a_4_**). A second nicotinic response was also found in neurons that exhibited a burst-firing action potential firing pattern (**b_1_**). Decay kinetics of this nicotinic response contained both fast and slow components (**b_2_**). The fast component was inhibited by 10 nM MLA (**b_3_**) and the remaining slow component was inhibited by 3 μM DHβE (**b_4_**), which suggests the presence of both homomeric α7 and heteromeric β2* nAChRs. A third nicotinic response was found in neurons that exhibited a regular-firing action potential firing pattern (**c_1_**). This nicotinic responses had slow rise and decay kinetics characteristic of heteromeric β2* nAChRs (**c_2_**). This was confirmed by the lack of effect of 10 nM MLA (**c_3_**) and the complete inhibition by 3 μM DHβE (**c_4_**). **d** Pie charts show the proportion of sampled neurons that exhibited each type of firing pattern and nicotinic response for male and female mice. These proportions were not statistically different between sexes (Chi-square test, *X*^2^ (2, *n* = 120) = 0.55, *p* = 0.76).

Previous studies have characterized the kinetics of nicotinic responses mediated by α7, α7β2 and β2* nAChRs^48,50^. We measured the kinetics of the three nAChR response types observed in this study by fitting current responses to an eight-term Fourier series equation (Fig. 2). Current amplitude (Fig. 2a) differed amongst response types (two-way ANOVA, *F*(2,112) = 16.29, *p* <0.0001) with α7 responses having a greater amplitude than α7/β2* and β2* responses in males and β2* responses in females (Bonferroni *post hoc* correction, each *p* < 0.01). The net charge (Fig. 2b) differed amongst response types (*F*(2,114) = 64.61, *p* <0.0001) with the β2* response having greater net charge than α7 and α7/β2* responses in both sexes (each *p* < 0.01). The time to the peak response (Fig. 2c) differed amongst response types (*F*(2,114) = 196.10, *p* <0.0001 with the β2* response having a greater time to peak than α7 and α7/β2* responses in both sexes (each *p* < 0.0001). Similarly, rise tau (Fig. 2d) differed amongst response types (*F*(2,114) = 84.42, *p* <0.0001 with the β2* response having a greater rise tau than α7 and α7/β2* responses in both sexes (each *p* < 0.0001). The first decay tau (Fig. 2e) differed amongst response types (*F*(2,114) = 418.80, *p* <0.0001) with the β2* response having a greater first decay tau than α7 and α7/β2* responses in both sexes (each *p* < 0.0001). This measure also differed between α7 and α7/β2* responses in females only (p < 0.01). The second decay tau (Fig. 2f) differed amongst response types (*F*(2,114) = 361.60, *p* <0.0001) and reliably differentiated the three response types, as each was different than the other in both sexes (each *p* < 0.0001). This measure is particularly sensitive to differentiating α7 and α7/β2* responses, as the second decay tau was less than 1000 ms for α7 responses and greater than 1000 ms for α7/β2* responses. These analyses demonstrate that whole-cell α7, α7/β2*, and β2* nicotinic responses have distinct activation and desensitization kinetics in juvenile mouse mPFC layer V pyramidal neurons.

**Fig. 2.**
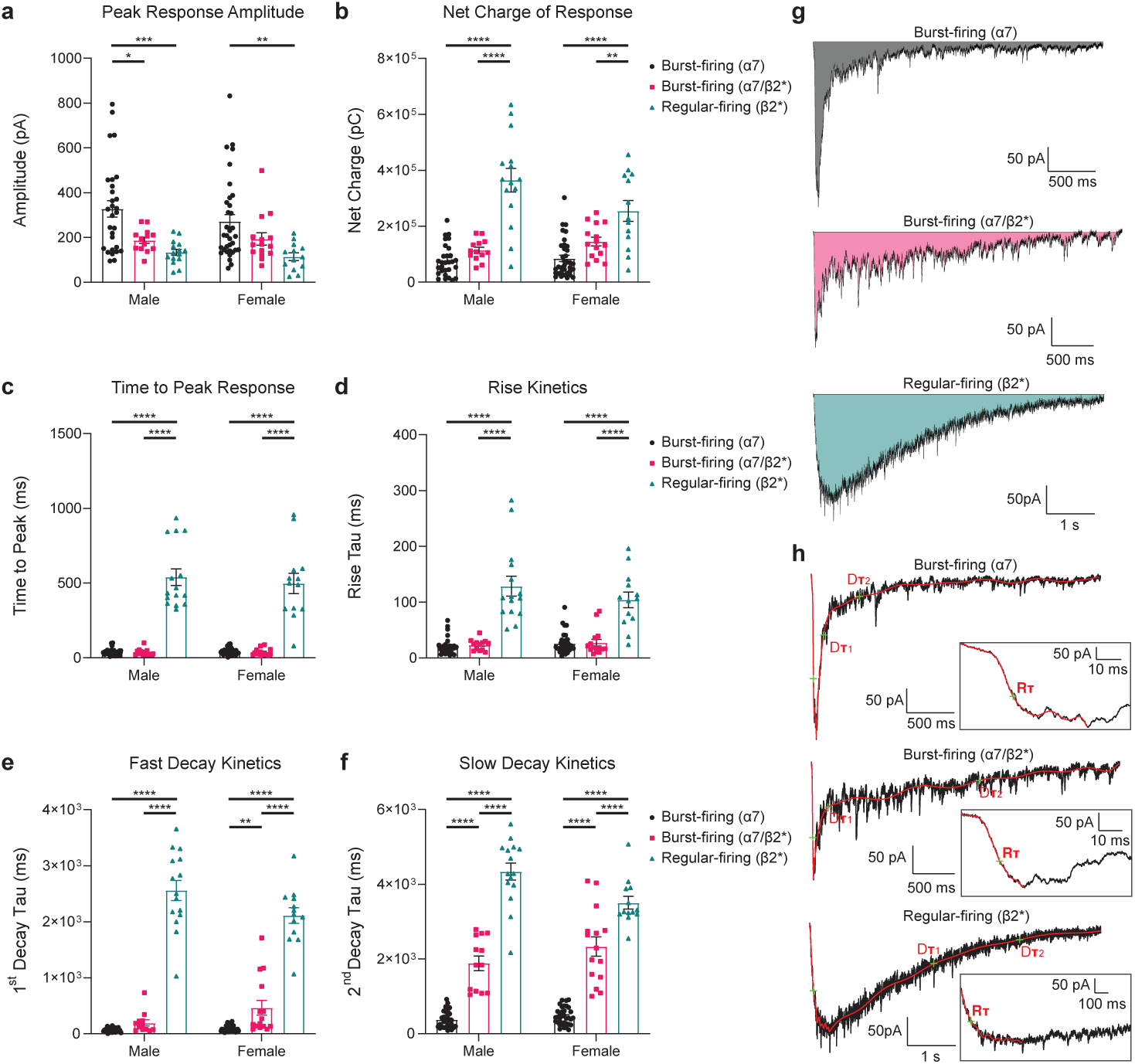
Characterization of nicotinic responses in prefrontal layer V pyramidal neurons. A detailed analysis of inward current response kinetics was performed for the three pharmacologically-identified nicotinic responses that were observed within each neuron subtype. Peak response amplitude (**a**) and net charge (**b**) showed no sex difference (two-way ANOVA, each *p* ≥ 0.2) but did show opposite effects of response type (each *p* < 0.0001). The time to peak (**c**) and rise tau (**d**) showed no sex difference (each *p* ≥ 0.4), but were both affected by response type (each *p* ≤ 0.0001) with β2* nAChR responses exhibiting the slowest response onset. The first decay tau (**e**) and second decay tau (**f**) showed no sex difference (each *p* ≥ 0.4), but were both affected by response type (each *p* ≤ 0.0001) with β2* nAChR responses exhibiting the slowest response decay amongst the response types. Of note, the second decay tau was significantly different between each the α7, α7/β2*, and β2* nAChR responses within each sex (Bonferroni *post hoc* correction, each comparison *p* < 0.0001). **g** One typical tracing is shown for each nicotinic response type with the solid area under the curve representing the net charge conducted. **h** The same typical tracings are shown with an eight-term Fourier series curve fitting to determine rise and decay kinetics. The rise tau (Rτ) is indicated by the green cross within each expanded box, while the first decay tau (Dτ_1_) and second decay tau (Dτ_2_) are indicated by green crosses within the full tracings. All data are shown for individual neurons and as mean +/-1 SEM. Sample sizes for males are: burst-firing (α7) = 29; burst-firing (α7/β2*) = 13; regular-firing (β2*) = 15, and for females are: burst-firing (α7) = 35; burst-firing (α7/β2*) = 15; regular-firing (β2*) = 13. Significance levels for Bonferroni *post hoc* corrections within each sex are \**p* < 0.05, \*\**p* < 0.01, \*\*\**p* < 0.001, and \*\*\*\**p* < 0.0001.

### Intrinsic electrophysiological properties of layer V pyramidal neuron subtypes

Having categorized the recorded pyramidal neurons into three subtypes based on action potential firing patterns and nicotinic response types, we next analyzed the intrinsic electrophysiological properties of each neuron subtype (Fig. 3). Two-way ANOVA demonstrated that resting membrane potential does not differ between neuron subtypes (*F*(2,114) = 2.33, *p* = 0.1) or sexes (*F*(1,114) = 0.25, *p* = 0.6). Input resistance (Fig. 3a) was affected by neuron subtype (*F*(2,114) = 29.26, *p* < 0.0001) with resistance being greater in RF (β2*) neurons than in each of the BF subtypes (Bonferroni *post hoc* correction, each *p* < 0.001). As expected, ISI ratio (Fig. 3b) was affected by neuron subtype (*F*(2,113) = 46.71, *p* < 0.0001) with the ISI ratio being greater in RF (β2*) neurons than in each of the BF subtypes (each *p* < 0.001). Rheobase (Fig 3c) showed an effect of neuron subtype (*F*(2,113) = 5.10), *p* = 0.008) although sag ratio (Fig 3d) did not *F*(2,114) = 2.67, *p* = 0.07). Additional analyses revealed differences in active electrophysiological properties between neuron subtypes. For example, within input/output experiments RF (β2*) neurons exhibited greater excitability and a noticeable depolarization block in both sexes (Fig. 3e-f). The magnitude of the post-firing afterhyperpolarization (AHP), which influences excitability, also differed between neuron subtypes. In both sexes, there was a significant interaction between the number of action potentials and AHP magnitude (Fig. 3g-h). Overall, these analyses demonstrate that the passive and active intrinsic electrophysiological properties differ between the layer V pyramidal neuron subtypes of this study, with the two BF subtypes being similar to each other and different from the RF (β2*) subtype.

**Fig. 3.**
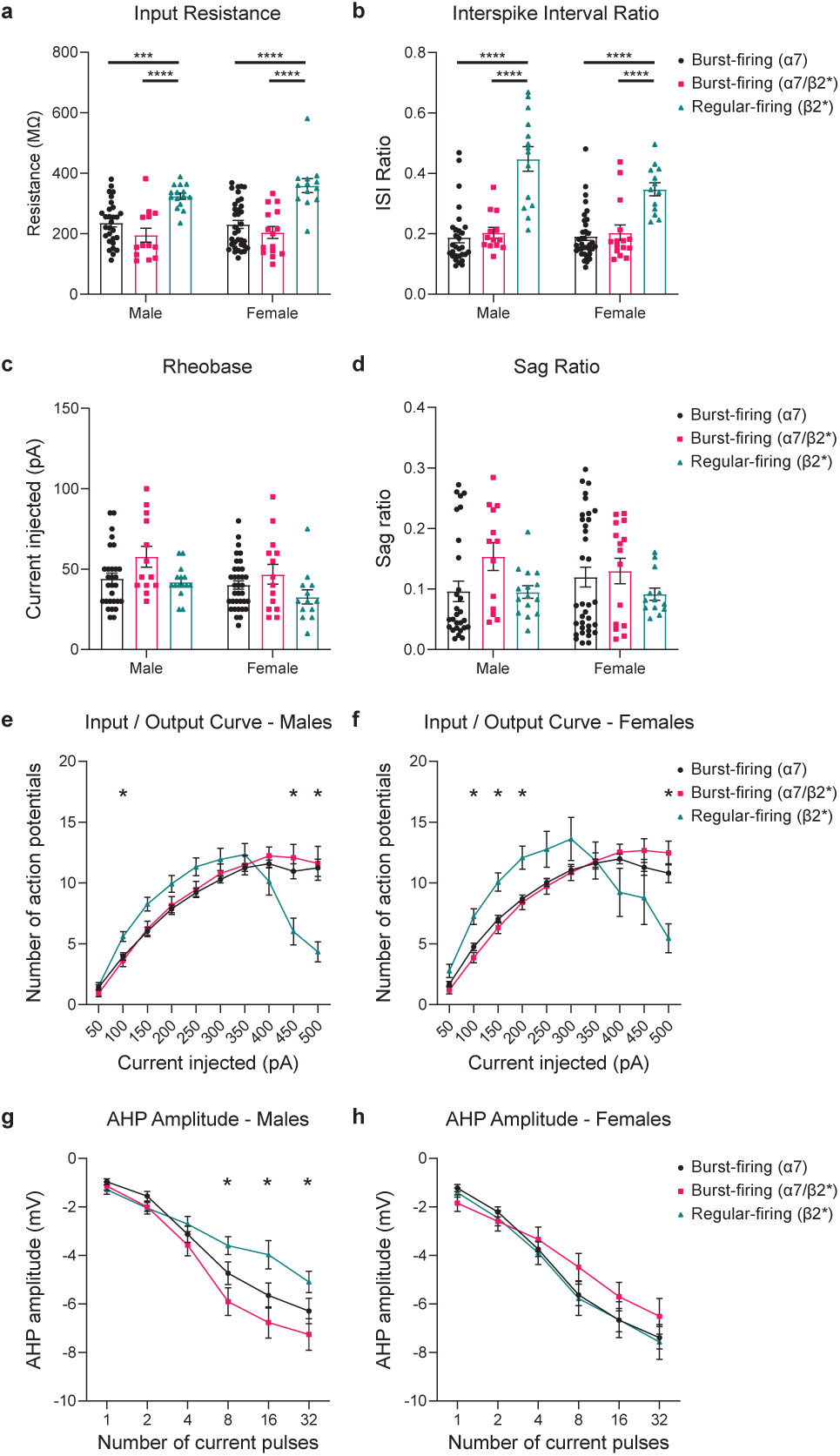
Characterization of intrinsic electrophysiological properties of prefrontal layer V pyramidal neurons. Input resistance (**a**) showed no sex difference (two-way ANOVA, *p* = 0.4), but was affected by neuron subtype (*p* <0.0001) with resistance being greatest in regular-firing (β2*) neurons (Bonferroni *post hoc* correction, all *p* ≤ 0.001). The same scenario was observed for interspike interval ratio (**b**), which showed no sex difference (*p* = 0.09), but was affected by neuron subtype (*p* < 0.0001) with the interspike interval ratio being greatest in regular-firing (β2*) neurons (all *p* ≤ 0.001). Rheobase (**c**) showed a sex difference (*p* = 0.02) and was affected by neuron subtype (*p* = 0.008). Sag ratio (**d**) showed no sex difference (*p* = 0.09) and was not affected by neuron subtype (*p* = 0.07). Input / output curves showed similar effects of neuron subtype within males (**e**) and females (**f**), where two-way ANOVA identified no overall effect of neuron subtype (males: *p* = 0.8; females: *p* = 0.2) but a significant interaction between effects of neuron subtype and the amount of current injected (*p* < 0.0001 for each sex). Firing output was different in regular-firing (β2*) neurons than in the other neuron subtypes where indicated (each *p* < 0.05). A similar scenario was observed for AHP amplitude in males (**g**) and females (**h**), with no overall effect of neuron subtype (males: *p* = 0.1; females *p* = 0.7) but a significant interaction between effects of neuron subtype and the number of current pulses (males: *p* < 0.0001; females: *p* = 0.02). AHP amplitude was lower in regular-firing (β2*) neurons than in burst-firing (α7/β2*) neurons for males only (as indicated; each *p* < 0.05). All data are shown for individual neurons and/or as mean +/-1 SEM. Sample sizes for males are: burst-firing (α7) = 29; burst-firing (α7/β2*) = 13; regular-firing (β2*) = 15, and for females are: burst-firing (α7) = 35; burst-firing (α7/β2*) = 15; regular-firing (β2*) n = 13. Significance levels for Bonferroni *post hoc* corrections within each sex are \**p* < 0.05, \*\**p* < 0.01, \*\*\**p* < 0.001, and \*\*\*\**p* < 0.0001.

### Excitatory presynaptic input to layer V pyramidal neuron subtypes

The mPFC receives excitatory input from cortical and subcortical sources. We measured the properties of spontaneous excitatory postsynaptic currents (EPSCs) received by the three layer V pyramidal neuron subtypes that were identified in this study. EPSC frequency (Fig. 4a) was affected by neuron subtype (two-way ANOVA, *F*(2,113) = 13.41, *p* < 0.0001), with frequency being lower in RF (β2*) neurons than BF (α7) neurons in both sexes (Bonferroni post hoc correction, *p* < 0.01). Similarly, both EPSC amplitude (Fig. 4b, F(2,112) = 10.31, p < 0.0001) and net charge (Fig. 4c, F(2,113) = 17.26, p < 0.0001) were affected by neuron subtype with values being lower in RF (β2*) neurons than in BF (α7) neurons in both sexes (all p < 0.01). There was no sex difference for EPSC frequency (F(1,113) = 0.35, p = 0.6) but there were sex differences for both EPSC amplitude (F(1,112) = 20.34, p < 0.0001) and net charge F(1,113) = 19.91, p < 0.0001) in that values were greater in males than females. These results suggest that excitatory synaptic signalling received within mPFC layer V differs depending on pyramidal neuron subtype.

**Fig. 4.**
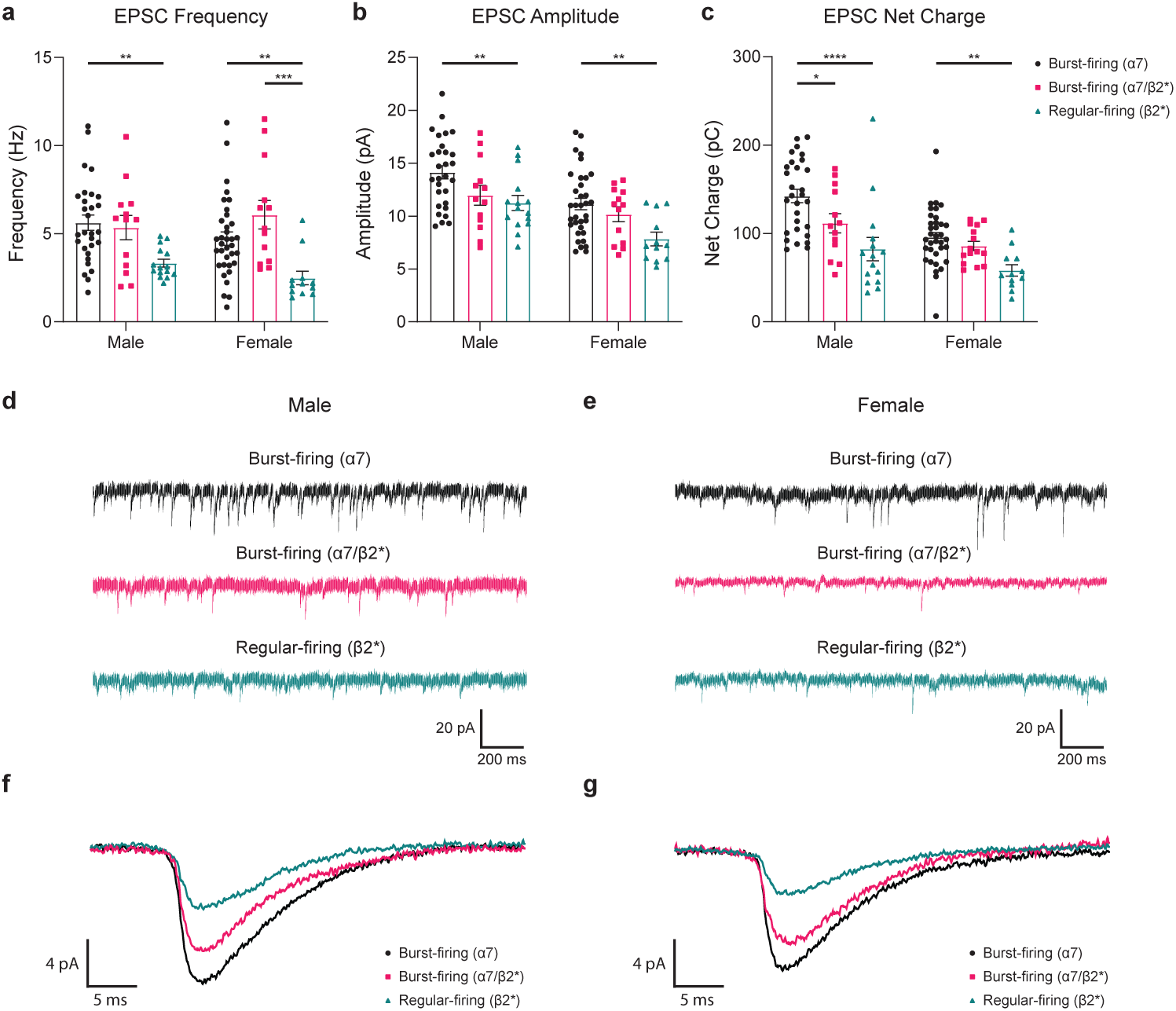
Characterization of spontaneous presynaptic input to prefrontal layer V pyramidal neurons. Spontaneous EPSC frequency (**a**) showed no sex difference (two-way ANOVA, *p* = 0.6), but was affected by neuron subtype (*p* < 0.0001) with frequency being lowest in regular-firing (β2*) neurons (Bonferroni *post hoc* correction, *p* < 0.01 or *p* < 0.001 as indicated). EPSC amplitude (**b**) showed a sex difference (*p* < 0.0001) and an effect of neuron subtype (*p* < 0.0001), with amplitude being lowest in regular-firing (β2*) neurons (*p* < 0.01 as indicated). Similarly, EPSC net charge showed a sex difference (*p* < 0.0001) and an effect of neuron subtype (*p* < 0.0001), with net charge being lowest in regular-firing (β2*) neurons (*p* < 0.05, *p* < 0.01, or *p* < 0.0001 as indicated). Typical voltage-clamp traces with EPSCs are shown for one pyramidal neuron from each subtype in males (**d**) and females (**e**). The average of 200 representative EPSCs is shown for pyramidal neurons from each subtype in males (**f**) and females (**g**). All data are shown for individual neurons and as mean +/-1 SEM. Sample sizes for males are: burst-firing (α7) = 29; burst-firing (α7/β2*) = 13; regular-firing (β2*) = 15, and for females are: burst-firing (α7) = 35; burst-firing (α7/β2*) = 15; regular-firing (β2*) n = 13. Significance levels for Bonferroni *post hoc* corrections within each sex are \**p* < 0.05, \*\**p* < 0.01, \*\*\**p* < 0.001, and \*\*\*\**p* < 0.0001.

### Correlation between electrophysiological properties and nicotinic responses

Correlation analyses were performed for all neurons within each sex, to assess relationships between eight intrinsic electrophysiological properties, six measures of nicotinic currents, and three measures of EPSCs. Fig. 5a. shows the resulting correlation matrix for each sex, with positive correlations marked green and negative correlations marked red. Multiple positive correlations were identified that could be expected based on group differences between pyramidal neuron subtypes. For example, input resistance (which was greatest in RF (β2*) neurons) correlated positively with interspike interval adaptation ratio and input/output excitability (which were both greatest in RF (β2*) neurons). Input resistance also correlated positively with measures of nicotinic response kinetics, which were all slower in RF (β2*) neurons. Correlations were identified between measures of nicotinic responses and EPSCs that were not expected. For example, measures of nicotinic response kinetics correlated negatively with EPSC frequency but correlated positively with EPSC amplitude and EPSC net charge.

**Fig. 5.**
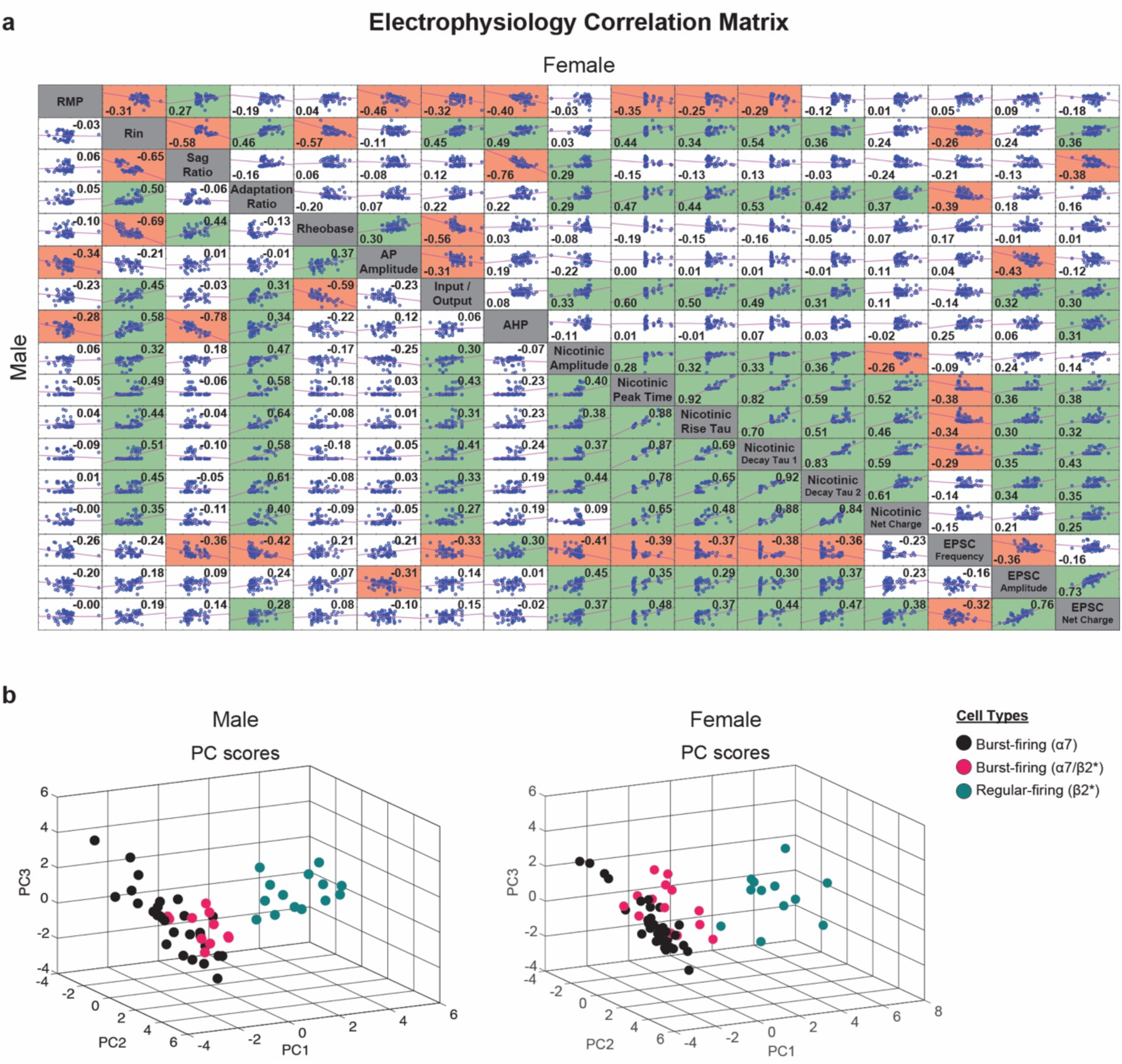
Correlation and principal component analyses for electrophysiological properties of prefrontal layer V pyramidal neurons. **a** Correlation matrix comprising eight intrinsic electrophysiological properties, six measures of nicotinic current responses, and three measures of spontaneous EPSCs. Male data is presented on the lower-left and female data is presented on the upper-right. For each comparison, the corresponding cell contains the Pearson correlation coefficient, a graphical representation of the correlation analysis, and the line of best fit. Cells with significant positive correlations are shown in green and cells with significant negative correlations are shown in red (each *p* < 0.05). **b** Principal component analysis included all measures from the correlation matrix. Three-dimension plots are shown for the first three principal components (PCs) that account for 65.9% of variability in males and 60.0% of variability in females. In both sexes, regular-firing (β2*) neurons form an unambiguous cluster separate from the other two burst-firing neuron subtypes. Sample sizes for males are: burst-firing (α7) = 29; burst-firing (α7/β2*) = 13; regular-firing (β2*) = 15, and for females are: burst-firing (α7) = 35; burst-firing (α7/β2*) = 15; regular-firing (β2*) n = 13.

We next assessed potential clustering of recorded neurons into groups using a principal component analysis, for which each of the 17 electrophysiological property, nicotinic response, and EPSC property measures used in the correlation analyses were treated as principal components (PCs). Neurons from male mice had four PCs and neurons from female mice had six PCs with eigenvalues greater than 1, which accounted for 74.2% and 83.2% of the total variance, respectively. A screeplot was used to confirm that a plateau was reached with these PCs and that the remaining PCs did not significantly contribute to group variance. The three PCs with the greatest contribution toward group variance accounted for 37.0%, 15.9%, and 13.1% of total variance in males and 33.4%, 15.1%, and 11.5% of the total variance in females. In both the sexes, the measures that largely contributed to the first PC are nicotinic peak response amplitude (Male: 0.13; Female: 0.14), rise tau (Male: 0.10; Female: 0.12), 1^st^ decay tau (Male: 0.13; Female: 0.14) and 2^nd^ decay tau (Male:0.12; Female: 0.10). Fig. 5b shows three-dimension plots of neurons positioned according to the three largest PCs. Here, neurons separated into two distinct clusters comprising the BF (α7 and α7/β2*) neurons or RF (β2*) neurons. For the BF neuron subtypes, BF (α7) neuron positions were more elongated along the PC2 and PC3 axes, whereas the BF (α7/β2*) neurons formed a tighter cluster within all three axes.

### Distinct morphologies of layer V pyramidal neuron subtypes

The morphology of mPFC layer V pyramidal neuron subtypes has been reported previously^31,32,34,44,45,51^. The morphology of neighbouring mPFC layer VI pyramidal neurons has been related previously with muscarinic^52^ and nicotinic^53^ responses, so we next sought to determine whether the morphology of mPFC layer V neurons in this study relates with their nicotinic response type. Sholl-type plots were prepared for a random subset of neurons within each sex that were labelled during electrophysiological recording and analysed based on dendrite length (Fig. 6a,b), diameter (Fig. 6c,d), and volume (Fig 6e,f).

**Fig. 6.**
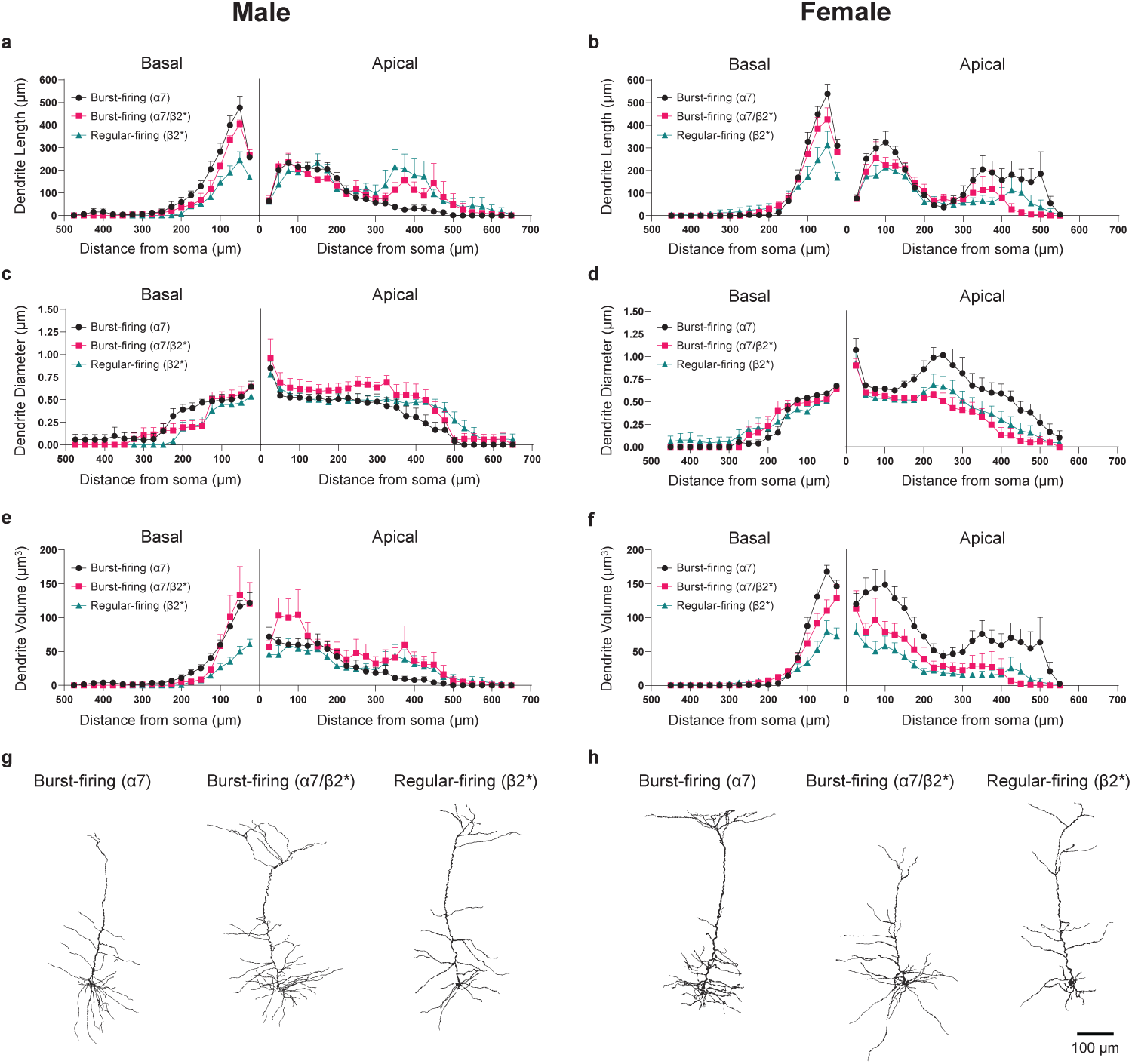
Dendrite morphology of prefrontal layer V pyramidal neurons. Sholl plots are shown for dendrite length, diameter, and volume as a function of distance from neuron soma. Data are shown for males on the left side of the figure and for females on the right side of the figure. For each plot, apical dendrite data is oriented to the right of zero, and basal dendrite data is oriented to the left of zero, with zero indicating the location of the soma. Apical dendrite length showed an effect of neuron subtype for both males (**a**) and females (**b**) (two-way ANOVA, each *p* < 0.0001). Burst-firing (α7) neurons showed the lowest apical length in males (Bonferroni *post hoc* correction, *p* ≤ 0.05) and the greatest apical length in females (*p* <0.0001). Basal dendrite length showed an effect of neuron subtype in both sexes (*p* ≤ 0.001) with the lowest basal length in regular-firing (β2*) neurons (*p* ≤ 0.05). Apical dendrite diameter showed an effect of neuron subtype in males (**c**) and females (**d**) (each *p* < 0.0001) whereas basal dendrite diameter showed an effect of neuron subtype in males only (*p* < 0.0001). Apical dendrite volume showed an effect of neuron subtype for both males (**e**) and females (**f**) (each *p* < 0.0001). Burst-firing (α7/β2*) neurons showed the greatest apical dendrite volume in males (*p* ≤ 0.001) whereas burst-firing (α7) neurons showed the greatest apical dendrite volume in females (*p* < 0.0001). Basal dendrite volume showed an effect of neuron subtype in both sexes (*p* ≤ 0.0001) with the lowest basal dendrite volume in regular-firing (β2*) neurons (*p* ≤ 0.001). Sample sizes for males are: burst-firing (α7) = 9; burst-firing (α7/β2*) = 6; regular-firing (β2*) = 7, and for females are: burst-firing (α7) = 11; burst-firing (α7/β2*) = 6; regular-firing (β2*) n = 8. Typical traces of reconstructed pyramidal neurons are shown for males (**g**) and females (**h**) below the Sholl plots for each sex. The scale bar represents 100 μm.

Apical dendrite length showed opposite effects of neuron subtype in each sex. In males, there was a significant effect of neuron subtype (two-way ANOVA, *F*(2,520) = 11.99, *p* < 0.0001) with BF (α7) neurons having a smaller length than the other two neuron subtypes (Bonferroni post hoc correction, each *p* < 0.01). In females, there was a significant effect of neuron subtype (*F*(2,506) = 16.52, *p* < 0.0001) with BF (α7) neurons having a greater length than the other two neuron subtypes (each *p* < 0.0001). Basal dendrite length showed similar effects of neuron subtype within males (*F*(2,380) = 41.55, *p* < 0.0001) and females (*F*(2,414) = 8.82, *p* = 0.0002). In both sexes, BF (α7) neurons had the greatest basal dendrite length and RF (β2*) neurons had the smallest basal dendrite length. All neuron subtypes were different from each other (each *p* < 0.05) except for the two BF subtypes within females.

Apical dendrite diameter showed opposite effects of neuron subtype in each sex. In males, there was a significant effect of neuron subtype (*F*(2,520) = 20.81, *p* < 0.0001) with BF (α7) neurons having a smaller diameter than the other two neuron subtypes (each *p* < 0.001). In females, there was a significant effect of neuron subtype (*F*(2,506) = 53.13, *p* < 0.0001) with BF (α7) neurons having a greater diameter than the other two neuron subtypes (each *p* < 0.0001). Basal dendrite diameter showed an effect of neuron subtype within males (*F*(2,380) = 16.65, *p* < 0.0001) with BF (α7) neurons having a greater diameter than the other two neuron subtypes (each *p* < 0.05). Basal dendrite diameter was not affected by neuron subtype in females (*F*(2,414) = 1.60, *p* = 0.2).

Apical dendrite volume showed opposite effects of neuron subtype in each sex. In males, there was a significant effect of neuron subtype (*F*(2,520) = 14.19, *p* < 0.0001) with BF (α7) neurons having a smaller volume than BF (α7/β2*) neurons (*p* < 0.0001). In females, there was a significant effect of neuron subtype (*F*(2,506) = 65.19, *p* < 0.0001) with BF (α7) neurons having a greater volume than the other two neuron subtypes (each *p* < 0.0001). Basal dendrite volume showed similar effects of neuron subtype within males (*F*(2,380) = 18.79, *p* < 0.0001) and females (*F*(2,414) = 25.46, *p* < 0.0001), with RF (β2*) neurons having a smaller basal dendrite volume than the other two neuron subtypes (each p < 0.001).

### Sex differences in the morphology of layer V pyramidal neuron subtypes

We performed a direct comparison of dendrite volume between male and female pyramidal neurons sampled from each subtype. The most striking sex difference was observed for BF (α7) neurons, where apical (two-way ANOVA, *F*(1,468) = 110.20, *p* < 0.0001) and basal (*F*(1,342) = 5.57, *p* = 0.02) dendrites had a greater volume in females (Fig. 7a). On the contrary, there was no sex difference for BF (α7/β2*) neuron apical (*F*(1,213) = 3.69, *p* = 0.06) or basal (*F*(1,228) = 0.03, *p* = 0.9) dendrites (Fig. 7b). There were subtle sex differences in RF (β2*) neurons, where apical dendrites had a greater volume in males (*F*(1,338) = 4.77, *p* = 0.03) and basal dendrites had a greater volume in females (*F*(1,247) = 11.82, *p* = 0.0007) (Fig. 7c). These results demonstrate that the morphology of certain mPFC layer V pyramidal neuron subtypes differs between the sexes during the juvenile period.

**Fig. 7.**
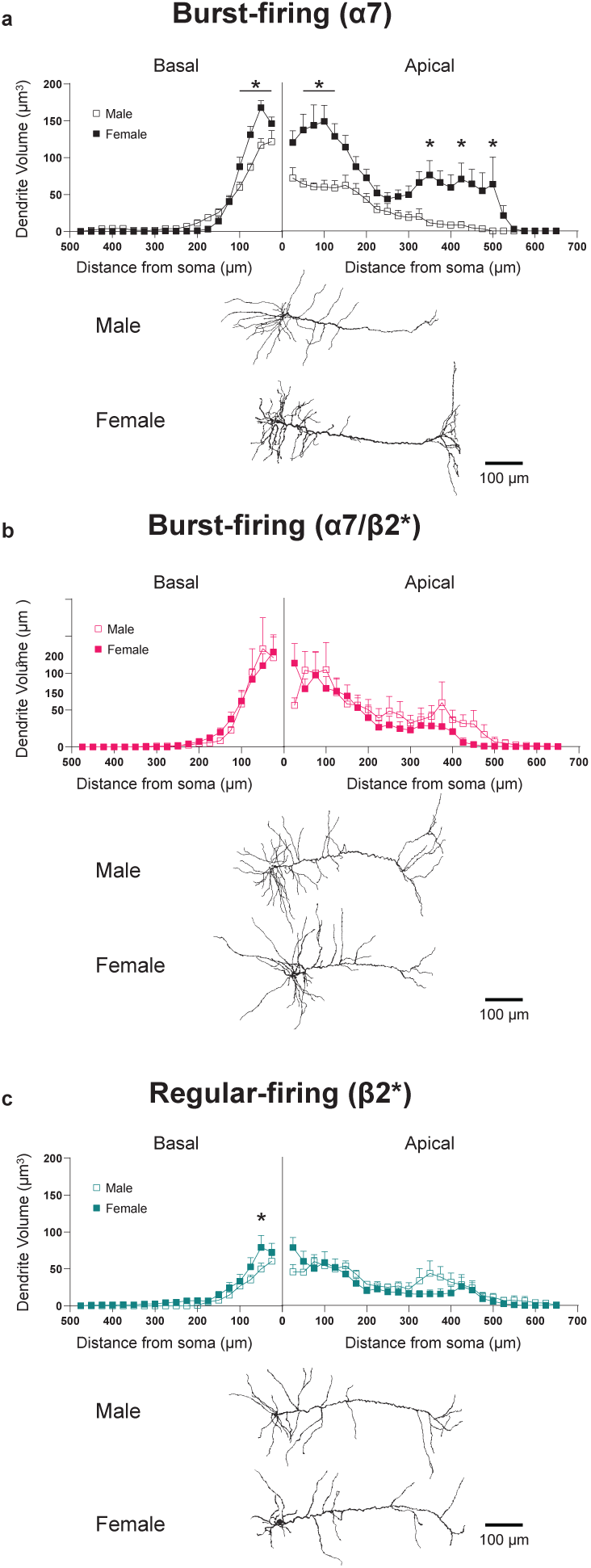
Sex differences in the morphology of prefrontal layer V pyramidal neurons. Sholl plots are shown comparing male and female dendrite volume for each of the pyramidal neuron subtypes. For each plot, apical dendrite data is oriented to the right of zero, and basal dendrite data is oriented to the left of zero, with zero indicating the location of the soma. **a** In burst-firing (α7) neurons, apical dendrite volume showed a sex difference (two-way ANOVA, *p* <0.0001) that depended on the distance from the soma (interaction effect, *p* = 0.005). Apical dendrites from females had a greater volume than those from males (Bonferroni *post hoc* correction, *p* < 0.05 as indicated). Basal dendrite volume also showed a sex difference (*p* = 0.02) that depended on the distance from the soma (*p* < 0.0001). **b** In burst-firing (α7/β2*) neurons, there was no sex difference for apical dendrite volume (*p* = 0.06) or for basal dendrite volume (*p* = 0.9). **c** In regular-firing (β2*) neurons, both apical dendrite volume (*p* = 0.03) and basal dendrite volume (*p* = 0.0007) showed a significant sex difference with no dependence on distance from the soma (both *p* = 0.6). Basal dendrite volume was greater in females, as indicated. Sample sizes for males are: burst-firing (α7) = 9; burst-firing (α7/β2*) = 6; regular-firing (β2*) = 7, and for females are: burst-firing (α7) = 11; burst-firing (α7/β2*) = 6; regular-firing (β2*) n = 8. All data are shown as mean +/- 1 SEM. \**p* < 0.05 for a significant sex difference following Bonferroni *post hoc* corrections. Typical traces of reconstructed neurons from each sex are shown below each Sholl plot. The scale bars represent 100 μm.

### Brain region projection targets of layer V pyramidal neuron subtypes

Previous work has demonstrated that electrophysiological and/or morphological properties of mPFC layer V pyramidal neurons are related with their axonal projection targets^32,44,45^. We next used a retrograde labelling approach to determine whether the mPFC layer V pyramidal neuron subtypes identified in this current study, based on action potential firing patterns and nicotinic response types, also exhibit distinct axonal projection targets. A retrograde tracer was injected into the contralateral mPFC, ipsilateral ventral striatum (nucleus accumbens), or ipsilateral ventromedial thalamus in mice of both sexes at P18 and labelled mPFC layer V pyramidal neurons were recorded for whole cell electrophysiology at P21. Neurons were recorded from 3-4 mice per projection target per sex. The results from this experiment confirm that each neuron subtype projects to a distinct cortical or sub-cortical target. All recorded cortico-cortical neurons projecting to the contralateral mPFC had electrophysiological properties consistent with the BF (α7) subtype (Fig. 8a), as they exhibited a BF action potential firing pattern and a nicotinic response with a fast onset and one-component fast decay. These properties were observed in 10/10 neurons from males and 10/10 neurons from females. All recorded cortico-striatal neurons projecting to the ipsilateral nucleus accumbens had electrophysiological properties consistent with the BF (α7/β2*) subtype (Fig. 8b), as they exhibited a BF action potential firing pattern and a nicotinic response with a fast onset and two-component fast plus slow decay.

**Fig. 8.**
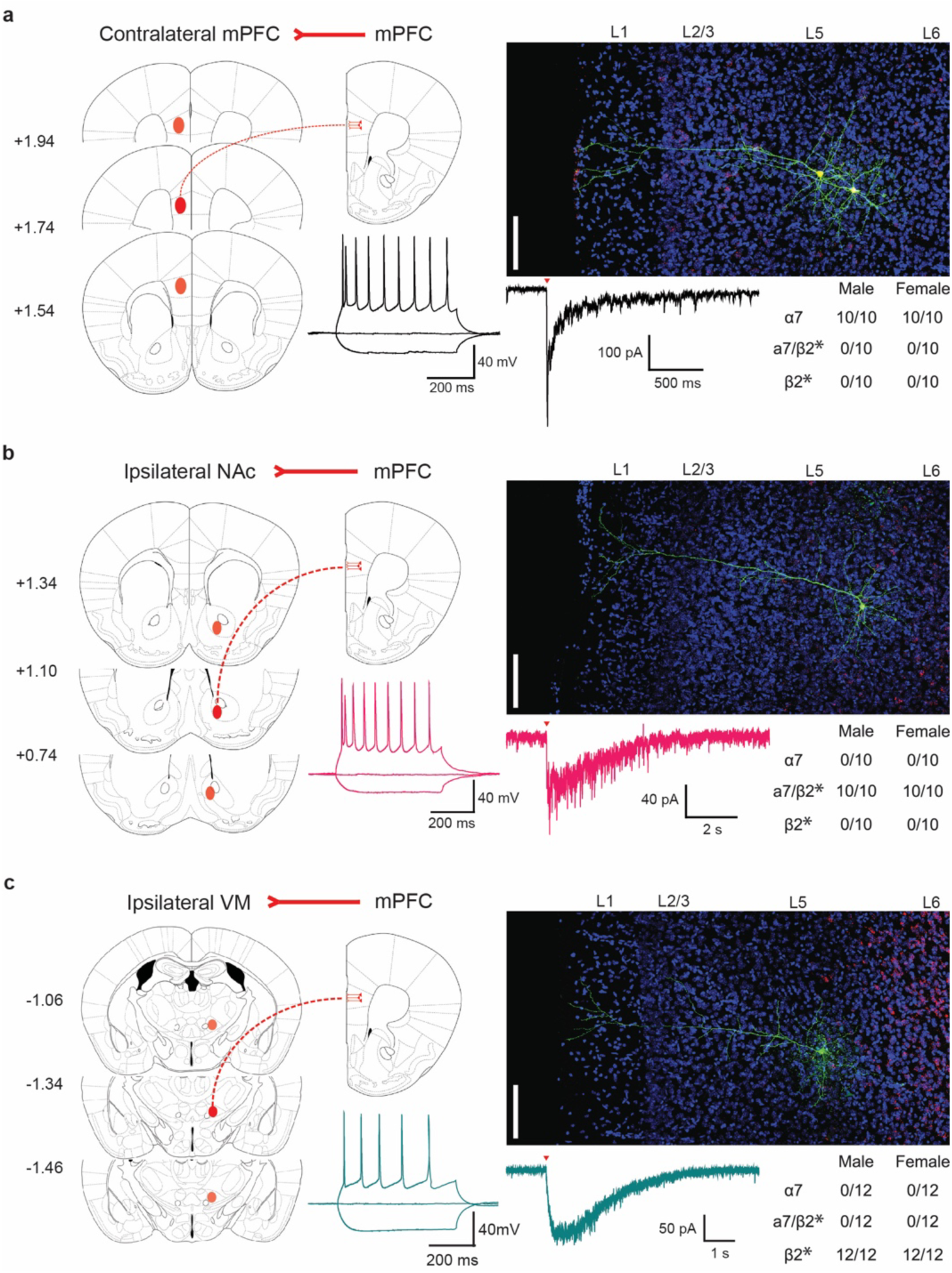
Retrograde labelling of prefrontal layer V pyramidal neurons. Prefrontal layer V pyramidal neurons were recorded after intracranial injection of retrograde tracer into the (**a**) contralateral mPFC, (**b**) ipsilateral nucleus accumbens, or (**c**) ipsilateral ventromedial thalamus. For each injection region, coronal diagrams relative to bregma show the injection site in red. Representative photomicrographs of the mPFC show DAPI-stained nuclei (blue), retrograde tracer (red), and layer V pyramidal neurons containing the retrograde tracer that were recorded for electrophysiology and filled with biocytin (green). For the contralateral mPFC, all retrograde-labelled neurons that were recorded showed a burst-firing response to 200 pA of positive current and an α7 nicotinic response to 1 mM ACh (indicated by the red arrowhead). No other nicotinic response was observed. For the ipsilateral nucleus accumbens, all retrograde-labelled neurons that were recorded showed a burst-firing response to 200 pA of positive current and an α7/β2* nicotinic response to 1 mM ACh. For the ipsilateral ventromedial thalamus, all retrograde-labelled neurons that were recorded showed a regular-firing response to 200 pA of positive current and a β2* nicotinic response to 1 mM ACh. The contingency table presented for each injection region shows the proportion of recorded neurons that showed each nicotinic response type. Scale bars on the photomicrographs represent 100 μm.

These properties were observed in 10/10 neurons from males and 10/10 neurons from females. In a separate set of animals, retrograde labelling showed that neurons projecting to the caudate/putamen, basolateral amygdala, and lateral hypothalamus also all had electrophysiological properties consistent with the BF (α7/β2*) subtype, suggesting that neurons from this subtype may project to multiple targets. All recorded cortico-thalamic neurons projecting to the ipsilateral ventromedial thalamus had electrophysiological properties consistent with the RF (β2*) subtype (Fig. 8c), as they exhibited a RF response to positive current injection and a nicotinic response with a slow onset and one-component slow decay. These properties were observed in 12/12 neurons from males and 12/12 neurons from females.

## Discussion

Layer V pyramidal neurons play a central role within mPFC cognitive circuits. They span the cortical column, integrate afferent signals from cortical and subcortical sources^54^, and send efferent signals to cortical and subcortical targets^55^. Their role to support cognitive functions is modulated by cholinergic neurotransmission that involves postsynaptic nicotinic responses^16,18,56^. Characterizing the manner by which nicotinic signalling modulates mPFC layer V neurons in juvenile age mice is important for understanding the role of cholinergic neurotransmission in the maturation and function of mPFC-associated circuits. In this present study, we report the following for mPFC layer V pyramidal neurons sampled from juvenile age mice: 1) Unique nAChR isoform-specific nicotinic responses are present in distinct subtypes of neurons; 2) these neuron subtypes exhibit distinct combinations of intrinsic electrophysiological and morphological properties, and axonal projection targets, which are largely consistent with previous reports; 3) these neuron subtypes exhibit distinct levels of spontaneous presynaptic excitatory input; and 4) strong correlations exist between measures of nicotinic responses and certain intrinsic electrophysiological properties. A graphical summary of neuron populations expressing α7 and β2* nAChRs within all layers of the mPFC is presented in Fig. 9, based on findings from this current study and previous reports.

**Fig. 9.**
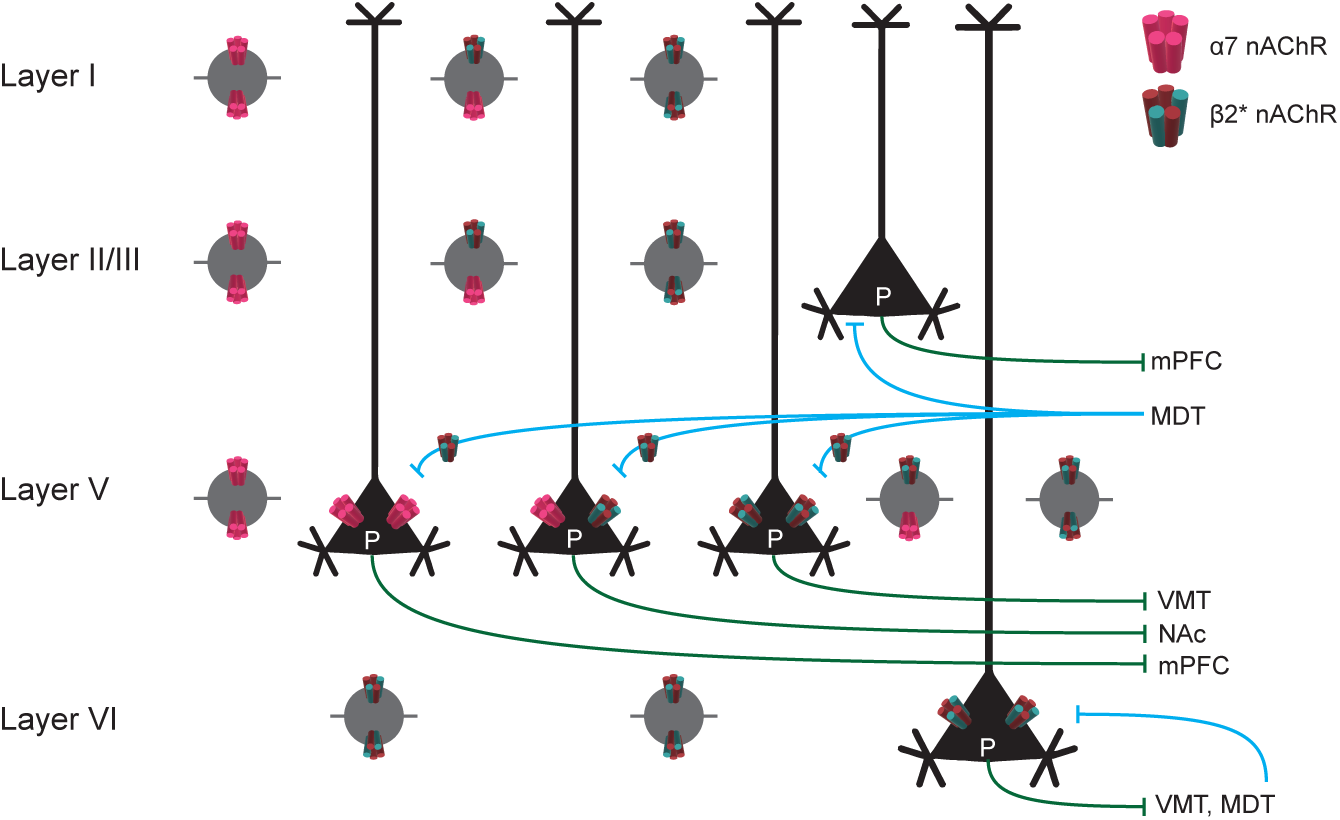
Graphical summary of nicotinic receptor localization within the prefrontal cortex. Based on previous literature and the findings from this current study, α7 and β2 subunit-containing nAChRs are expressed at distinct sites within the juvenile rodent medial prefrontal cortex. The majority of pyramidal neurons (P) in layers II/III do not appear to express nAChRs. Pyramidal neurons in layer V may be categorized into three subtypes based on this current study that express α7 and/or β2* nAChRs. Pyramidal neurons in layer VI express β2* nAChRs only. Interneurons in layers I, II/III, and V express α7 and/or β2* nAChRs, whereas interneurons in layer VI express β2* nAChRs only. Glutamatergic inputs from the mediodorsal thalamus to layer V pyramidal neurons also express β2* nAChRs. Glutamatergic inputs to the mPFC are shown in blue and outputs to specified projection targets are shown in green. Not all input or output pathways from the mPFC are shown. mPFC, medial prefrontal cortex; MDT, mediodorsal thalamus; VMT, ventrolateral thalamus; NAc, nucleus accumbens.

The functional presence of nAChR isoforms has been characterized previously for specific neuron types within each layer of the rodent mPFC^16,17,57,58^. That work shows α7 nAChRs to be present on fast spiking and non-fast spiking interneurons within layers I, II/III, and V, on a minority of pyramidal neurons within layer II/III, and on the majority of pyramidal neurons within layer V ^16,57,58^. It also shows β2* nAChRs to be present on thalamocortical glutamatergic terminals within layer V^59^, fast spiking interneurons within layer V^17^, non-fast spiking interneurons within all layers^16,58^, and on pyramidal neurons within layer VI only^16,53,60^. Our finding that nicotinic responses mediated by β2 subunit-containing nAChRs are present in approximately half of recorded BF neurons (α7/β2* nicotinic responses) and all of recorded RF neurons (β2* nicotinic responses) is not consistent with previous reports described above for mPFC layer V. A possible explanation for this inconsistency could be that we had actually recorded α7, α7/β2*, and β2* nicotinic responses in non-fast spiking interneurons as reported previously for this layer^16^. However, we are confident that we recorded from pyramidal neurons only, based on their characteristic action potential firing patterns, prominent apical dendrite morphology, and retrograde-labelled projections to known callosal and subcortical targets.

Differences in study design or recording conditions could underlie differences between our findings and previous reports. The work described above was performed using mice and rats of approximately the same developmental age as those in our study but was performed at a greater recording temperature than our study. Although recording temperature can transform action potential firing patterns between BF and RF for pyramidal neurons^61^, we have found that temperature does not influence apparent nAChR isoform-specific response types within the pyramidal neurons of our study. Another potential difference between our study and previous research is the mPFC sub-region from which neurons were sampled (i.e., the prelimbic or infralimbic regions). Neurons in our study were sampled from the prelimbic region only, whereas most previous reports do not state the sub-region from which neurons were sampled. There may be sex differences in nAChR expression or function in mPFC layer V, although we did not observe this for most measures of nAChR function in our study. Previous studies were performed in males only, combined males and females, or did not report the sex of experimental animals. Consistent with our findings, it should be noted that β2* nicotinic responses have been observed in layer V pyramidal neurons located within the rat somatosensory^18^ and mouse auditory^62^ cortices. However, details of these studies suggest that the mapping of isoform-specific nicotinic response types to specific pyramidal neuron subtypes may differ between the mPFC and primary sensorimotor cortices. For example, both α7 and β2* responses were observed in RF neurons of the somatosensory cortex^18^, and β2* responses were observed in both cortico-cortical and cortico-collicular neurons of the auditory cortex^62^.

Beyond the broad observation that nicotinic responses mediated by multiple nAChR isoforms are present in mPFC layer V pyramidal neurons, we show further that each unique nicotinic response type is present in a distinct neuron subtype. The neuron subtypes that we identified exhibit properties that are largely consistent with those described in previous reports for the mPFC. Based on axonal projection targets, IT neurons projecting within the contralateral cortex or to the striatum tend to show a BF action potential firing pattern and have a thinner apical dendrite tuft, compared with PT neurons projecting to subcortical regions that tend to show an RF action potential firing pattern and have a wider apical dendrite tuft^32,38,44,45,63^. Also consistent with our findings, Wang and colleagues^34^ found non-accommodating RF neurons with complex/wide apical dendrite tufts and accommodating BF neurons with simple/narrow apical dendrite tufts. This previous work largely matches the properties of neuron subtypes identified for male mice in our study, and only differs from female mice in our study with regard to their wide apical dendrite tuft for BF (α7) neurons. van Aerde and Feldmeyer^31^ identified neuron subtypes in rat mPFC based on action potential firing pattern and apical dendrite morphology, but observed the BF doublet only in a small fraction of neurons, and instead sub-divided RF neurons into adapting/non-adapting groups. Multiple aspects of experimental design differed between that study and ours, including species (rat), sex (both combined), age (P24-46) and recording temperature (30°C). The physiology and morphology of pyramidal neurons are well known to change during postnatal maturation into adulthood. Although the proportion of IB and RF layer V pyramidal neurons shifts between P18-22 and adulthood for the rat sensorimotor cortex^23^, the action potential firing patterns and apical dendrite morphology do not appear to differ between P13-16 and P26-30 in the mouse mPFC^51^. Shifting from room temperature to physiological temperature can shift the action potential firing pattern for layer V pyramidal neurons from IB to RF^61^, so it is possible that our recordings at room temperature do not reflect the neurons’ true action potential firing patterns *in vivo*. Notwithstanding this, our experimental design allowed us to identify the distinct neuron subtypes that were then characterized through their distinct nicotinic response types, morphology, and projection targets. Properties of the layer V pyramidal neuron subtypes themselves seem to differ between the mPFC and primary sensorimotor cortices, where BF neurons typically have thick apical dendrite tufts and project to PT targets whereas RF neurons typically have thin apical dendrite tufts and project to IT targets (see Introduction).

The most striking sex difference observed in this study was the greater apical and basal dendrite length and volume measured in BF (α7) neurons from female mice (Fig. 7a), which is not consistent with the greater EPSC amplitude and net charge measured in BF (α7) neurons from male mice (Fig. 4b, c). Other measures of intrinsic electrophysiological properties that may reflect neuron size (e.g. input resistance and rheobase) did not differ between sexes for this neuron type. These data suggest that neurons from male mice may be coupled with presynaptic glutamatergic input more efficiently in males than in females, or that the quantal release of glutamate is greater in males than in females, at this age. The only sex difference that we observed for nicotinic responses was the greater net charge of β2* nicotinic responses in neurons from male mice, which resulted from the slower response decay kinetics in this sex. We observed no sex difference in α7 or α7/β2* nicotinic responses. In neighbouring layer VI pyramidal neurons from the juvenile mouse mPFC, the β2* nicotinic response amplitude was reported to be greater in males than in females^60^.

We observed that nicotinic response types are related with apical dendrite morphology in a sex-dependent manner for mPFC layer V pyramidal neurons during the juvenile period of postnatal development. Nicotinic α7 responses map onto narrow tufted neurons in males and wide tufted neurons in females, whereas α7/β2* and β2* responses map onto neurons that are relatively wide tufted in males and narrow tufted in females. Cholinergic^64^, nicotinic^53^, and muscarinic^52^ signalling has been related with apical dendrite morphology for pyramidal neurons within other layers of the mPFC. It cannot be determined based on our data whether the nAChR isoform-specific nicotinic responses play a role to shape the dendrite morphology of the layer V pyramidal neuron subtypes identified in this study, although this would be an interesting avenue for future research.

Pyramidal neurons within layer V of the mPFC provide the primary output signal from this region and project to a wide variety of targets within limbic-cognitive circuits, including the mPFC itself and other regions of the cerebral cortex, ventral striatum (nucleus accumbens), dorsal striatum (caudate/putamen), thalamus, lateral hypothalamus, basolateral amygdala, ventral tegmental area, and brainstem/spinal cord^55,65,66^. Pyramidal neuron subtypes likely have specific roles to support cognitive functions, depending on their projection targets, intrinsic electrophysiological properties, and postsynaptic neurotransmitter receptors. In dorsal cortical regions, subtypes of IT and PT neurons show unique activity during, and contributions toward, an auditory decision task, with PT neurons making the greatest contribution^67,68^. In the mPFC, layer V pyramidal neurons projecting to the dorsal striatum contribute toward cognitive flexibility^69^ and inhibitory control^70^, while those projecting to the mediodorsal thalamus contribute toward attention and inhibitory control depending on the specific target within that brain region^70^. Regarding action potential firing patterns, it is the output from prefrontal BF neurons that is thought to synchronize this region during attention tasks^71,72^. Attention performance likely is supported by phasic^5^ and tonic^4^ ACh neurotransmission. Since nAChR activation and desensitization kinetics differ between nAChR isoforms, with faster activation and desensitization in α7 nAChRs than β2* nAChRs (reviewed in Papke and Lindstrom^73^), the presence of unique isoform-specific nicotinic responses in distinct pyramidal neuron subtypes likely influences their cellular response to ACh thereby shaping their distinct contributions toward mPFC-dependent and ACh-supported cognitive tasks.

## Methods

### Experimental animals

All experiments were performed using CD1-strain mice in accordance with the principles and guidelines of the Canadian Council on Animal Care, and under protocols that were approved by the University of Guelph Animal Care Committee. Pregnant mice were purchased from Charles River Canada (Saint-Constant, Quebec, Canada) and used to establish a local breeding colony. Mice were housed at an ambient temperature of 21-24°C with lights maintained on a 12-hour cycle (lights on at 8:00 pm) and were provided *ad libitum* access to food and water.

### Whole-cell electrophysiology in acute brain slices

Male and female mice between postnatal day (P) 15-20 were anesthetized using isoflurane and killed by decapitation. Whole brains were rapidly removed under 4°C oxygenated sucrose artificial cerebrospinal fluid (sACSF) (in mM: 245 sucrose, 10 D-Glucose, 26 NaHCO_3_, 2 CaCl_2_, 2 MgSO_4_, 3 KCl, and 1.25 NaH_2_PO_4_, pH 7.4). Coronal slices containing the mPFC from approximately Bregma +1.98 mm to +1.34 mm^74^ were made at 400 μm thickness using a Leica VT1200 vibrating microtome (Leica Microsystems Inc., Concord, Ontario, Canada). Slices were transferred to a recovery chamber containing oxygenated regular ACSF (in mM; 128 NaCl, 10 D-glucose, 26 NaH_2_CO_3_, 2 CaCl_2_, 2 MgSO_4_, 3 KCl, and 1.25 NaH_2_PO_4_, pH 7.4) and maintained at 30°C for at least two hours prior to electrophysiology recordings.

Slices were transferred to a recording chamber (Warner Instruments, Hamden, Connecticut, USA), superfused with fresh oxygenated ACSF at room temperature, and visualized by infrared differential interference contrast microscopy using an Axioskop FS2 microscope (Carl Zeiss Canada, Toronto, Ontario, Canada). Layer V pyramidal neurons were sampled for recording randomly within layer V of the prelimbic region, based on their location approximately 300-550 μm from the pial surface^16^, large soma size, pyramidal shape, and prominent apical dendrite projecting toward the pial surface. Whole-cell electrophysiological recordings were conducted using borosilicate glass electrodes having an internal resistance of 3-5 MΩ (Sutter Instrument Company, Novato, California, USA, catalogue number BF150-110-10). The electrode solution contained (in mM): 120 K-gluconate, 5 KCl, 2 MgCl_2_, 4 K_2_-ATP, 0.4 Na_2_-GTP, 10 Na_2_-phosphocreatine, 10 HEPES buffer (pH 7.3). Recordings were made using a Multiclamp 700B amplifier (Molecular Devices, San Jose, California, USA), with signals acquired at 20 kHz and low-pass filtered at 2 kHz using a Digidata 1440A digitizer (Molecular Devices). The liquid junction potential was corrected at the time of recording.

### Measurement of intrinsic electrophysiological properties

Neuron intrinsic electrophysiological properties were first measured in current clamp mode. Resting membrane potential was measured shortly after the whole-cell configuration was achieved with no current injection. Input resistance was calculated based on the change in membrane potential from rest in response to 500 ms of -100 pA current. Inter-spike interval (ISI) ratio was used to assess spike frequency adaptation and was calculated as the ratio of durations between the first two action potentials and last two action potentials generated in response to 500 ms of +200 pA current. Sag ratio was calculated as the ratio between the lowest membrane potential and the steady-state membrane potential generated in response to 500 ms of -100 pA current. Rheobase was measured by injecting 500 ms of positive current that increased in magnitude by 5 pA with each step until a single action potential was elicited. Action potential amplitude was measured at rheobase as the difference between the threshold potential and the peak potential. Postburst afterhyperpolarization (AHP) responses were measured as the difference between the resting membrane potential and the lowest membrane potential attained after injection of 1, 2, 4, 8, 16, and 32 current pulses (2 ms, 2 nA, at 50 Hz). Input/output curves were generated by measuring the number of action potentials elicited in response to 500 ms positive current steps that increased in 50 pA increments.

### Measurement of spontaneous excitatory postsynaptic currents

Spontaneous EPSCs were recorded in voltage-clamp mode at a holding potential of −75 mV for a 30s period. Tracings were analyzed offline using a template search in Clampfit software (version 10, Molecular Devices). Events that were <5 or >100 pA were not included in the analysis. EPSC frequency, amplitude, and net charge were measured.

### Measurement of postsynaptic nicotinic currents

Postsynaptic nicotinic currents were measured in voltage-clamp mode at -75 mV in the presence of 200 nM atropine to inhibit muscarinic receptors. 1 mM ACh was prepared in ACSF within a borosilicate glass application pipette having a resistance of 3-5 MΩ and pressure applied at 10 psi for 100 ms using a Picospritzer II (General Valve, Fairfield, New Jersey, USA). Relative to the soma, the application pipette was positioned 20-50 μm upstream and at a 45 degree angle from the flow of ACSF within the recording chamber. Initial experiments in Fig. 1 were performed in the presence of 10 nM MLA citrate (Tocris / Bio-Techne Canada, Toronto, Ontario, Canada) and/or 3 μM DHβE hydrobromide (Tocris) to determine the contribution of α7 and β2 subunit-containing nAChRs, respectively. Pressure application of ACSF alone did not alter the holding currents in these experiments.

Nicotinic currents were measured using a custom-written program in MATLAB (version 9.7, MathWorks, Natick, Maine, USA). Current amplitude, net charge, response time, rise and decay kinetics were used to characterized nicotinic currents mediated by α7, α7/β2*, and β2* responses (Fig. 2). Rise and decay kinetics were analyzed separately using nonlinear least-squares curve fitting via an eight-term Fourier series equation. Nicotinic currents mediated by α7 and β2* responses were distinguishable based on rise and decay kinetics, which were significantly more rapid for α7 responses than for β2* responses. In the subset of neurons that exhibited nicotinic currents mediated by α7/β2* responses, the second decay tau (Dτ_2_) threshold of 930 ms was used to distinguish between currents mediated by α7 and α7/β2* responses.

### Correlation matrix and principal component analysis

A total of eight measures of intrinsic electrophysiological properties, six measures of nicotinic currents, and three measures of EPSCs were used to generate a correlation matrix and perform principal component analysis within each sex (Fig. 5). The Pearson correlation coefficient and its associated *p* value were calculated for each possible pair of measures for all neurons irrespective of their nicotinic current response type. Principal component analysis was used to visualize the distribution of neuron subtypes (categorized based on nicotinic responses) in multidimensional space. The Kaiser-Gutman criterion was used to decide the initial number of PCs. Only PCs with an eigenvalue equal to or greater than one were selected for subsequent analysis. The selection of significant PCs was confirmed using a scree plot of all PCs. Significant PCs were then used to produce a three-dimension scatterplot matrix.

### Visualization and measurement of neuron morphology

During electrophysiological recordings, a subset of neurons exhibiting each type of nicotinic response were labelled with 0.3% (wt/vol) biocytin (Tocris) in the electrode solution. Brain slices were fixed in 4% (wt/vol) paraformaldehyde in 100 mM phosphate buffer (pH 7.5) for up to 3 weeks at 4°C. Slices were washed in Tris-buffered saline (TBS, 100 mM Tris and 150 mM NaCl, pH 7.5) 3 x 10 min at room temperature and incubated with 0.5% (vol/vol) H_2_O_2_ and 0.25% (vol/vol) Triton X-100 in TBS for 15 min at room temperature. Slices were washed in TBS 3 x 10 min at room temperature and incubated with 0.25% (vol/vol) VECTASTAIN Elite reagent A (avidin), 0.25% (vol/vol) VECTASTAIN Elite reagent B (biotinylated horseradish peroxidase), and 0.25% (vol/vol) Triton X-100 in TBS for 24 hours at room temperature (Vector Labs, Burlingame, California, USA). Slices were washed in TBS 3 x 10 min at room temperature and incubated with 0.05% (wt/vol) 3,3′-diaminobenzidine tetrahydrochloride hydrate (DAB), 0.04% (wt/vol) nickel chloride hexahydrate, and 0.25% (vol/vol) Triton X-100 in TBS for 5 min at room temperature. Slices were then incubated in this same DAB/nickel/Triton X-100 solution that also contained 0.001% (vol/vol) H_2_O_2_ for 15 minutes at room temperature. The reaction was stopped by washing slices 3 x 10 min in TBS at room temperature, after which slices were mounted onto microscope slides and allowed to air dry. Slides were dehydrated through ethanol gradients and coverslipped using Permount (Fisher Scientific, Ottawa, Ontario, Canada).

Individuals involved in neuron imaging and analysis were blinded to the experimental groups. Brightfield image stacks containing labelled neurons were captured using an Olympus BX53 upright microscope equipped with an Olympus UPlanSApo 30x / 1.05 N.A. silicone-immersion objective (Olympus Canada Inc., Richmond Hill, Ontario, Canada), which was controlled by Neurolucida software (MBF Bioscience, Williston, Vermont, USA). Labelled pyramidal neurons were reconstructed and measured only if they contained one prominent apical dendrite that projected toward the pial surface and all dendrites were fully contained within the slice. Neurons were reconstructed in three dimensions using Neuromantic software^75^ and their morphology was measured using Neurolucida Explorer software (MBF Bioscience). Modified Sholl analyses were conducted separately on apical and basal dendrites, in which the length, diameter, and volume of dendrites were measured between concentric spheres radiating from the soma in 25 μm increments.

### Retrograde labelling

Male and female mice underwent retrograde labelling surgery on P18. Three-to-four mice were used for each sex and labelling region. Carprofen was administered s.c. to the back at 20 mg/kg one hour before surgery. Mice were anaesthetised using isoflourane and placed in a stereotaxic apparatus (David Kopf Instruments, Tujunga, California, USA). The scalp was shaved and anesthetized with the s.c. administration of 50 µL of a local anesthetic cocktail containing 0.67% lidocaine and 0.17% bupivacaine, made from stock solutions in a volumetric ratio of 3:6:9 lidocaine:bupivacaine:saline. The scalp was cut along the midline and a small hole was drilled in the skull above the labelling region of interest. Relative to bregma, coordinates for each labelling region were: mPFC: AP: 1.8 mm, ML: -0.35 or 0.35 mm, DV: -2.6 mm, nucleus accumbens: AP: 1.1 mm, ML: -2.6 or 2.6 mm, DV -3.9 mm; 15° angle from vertical, and ventromedial thalamus: AP: -1.3 mm, ML: -0.75 or 0.75 mm, DV: -4.2 mm. A custom microinjector delivered 0.3 μL of rhodamine-labelled RetroBeads (0.5x dilution; LumaFluor Inc., Durham, North Carolina, USA) into the labelling region at a rate of 0.1 μL/min using infusion pump (model PHD 2000; Harvard Apparatus, Holliston, Maine, USA). Microinjectors were left in place for an additional 10 minutes, after which they were removed slowly and exposed skull was covered with dental cement. Mice were returned to their home cage for at least 48 h and killed for electrophysiology experiments at P20-21 as described above. Coronal brain sections containing the mPFC were visualized using widefield fluorescence and retrograde-labelled pyramidal neurons were assessed for physiology and morphology using whole-cell electrophysiology with biocytin labelling.

### Confocal Microscopy

After recording of retrograde-labelled neurons, brain slices were fixed in 4% (wt/vol) paraformaldehyde in 100 mM phosphate buffer overnight at 4°C. Slices were washed in TBS 3 x 10 min at room temperature and incubated in blocking buffer comprising 10% (wt/vol) bovine serum albumin (BSA) and 0.25% (vol/vol) Triton X-100 in TBS for two hours at room temperature. Slices were washed in TBS 3 x 10 min at room temperature and incubated in a solution containing 3% (wt/vol) BSA, 0.25% (vol/vol) Triton X-100, and 4 µg/mL streptavidin conjugated with Alex Fluor 488 (Fisher Scientific) in TBS overnight at 4°C. Slices were washed in TBS 3 x 10 min at room temperature and incubated with 1 μg/mL 4’,6-diamidino-2-phenylindole (DAPI) in TBS for two hours at room temperature. Slices were washed in TBS 3 x 10 min at room temperature and mounted onto microscope slides and coverslipped using an aqueous mounting medium.

Processed slices were imaged using an Olympus Fluoview FV1200 laser-scanning confocal system connected with an inverted Olympus IX83 microscope and controlled using Olympus Fluoview software (FV10-ASW version 4.0; Olympus). A helium neon laser tuned to 405 nm (DAPI), 488 nm (retrograde labelled neuron), or 543 nm (RetroBeads) and an Olympus UPlanSApo 30X, 1.05 NA silicone oil-immersion objective were used to acquire three-channel image stacks which, when stitched together in three dimensions, contained entire retrograde labelled neurons.

## Statistical Analysis

Electrophysiological recordings were analyzed using Clampfit software and custom scripts written for MATLAB software. Statistical analyses were conducted using GraphPad Prism software (version 9, GraphPad Software, Boston, Massachusetts, USA). Data were analysed using the Chi-square test, two-way ANOVA with the Bonferroni *post hoc* correction, or Pearson correlation, as indicated. Data are presented for individual neurons and as group means ± 1 SEM. For all figures, statistical significance is indicated as \**p* < 0.05, \*\**p* < 0.01, *** *p*< 0.001, and ****p < 0.0001. The sample size for each group is listed in the respective figure legend.

## Data Availability

All data that were generated and presented in this manuscript are available from the corresponding author (C.D.C.B.) upon reasonable request. Neuron reconstructions from this study will be submitted to NeuroMorpho.org.

## Acknowledgements

Funding for this work was provided by Discovery Grants from the Natural Sciences and Engineering Council of Canada (NSERC) to C.D.C.B. (RGPIN-2019-04989) and E.C. (RGPIN-2018-04699), and by a Canada Foundation for Innovation (CFI) grant to C.D.C.B. (30381). We thank Dr. Matthew Vickaryous at the University of Guelph for the use of a fluorescence filter cube.

## Author contributions

A.V.P. and C.D.C.B. conceived and designed the study. A.V.P., A.N., and P.P. performed experiments. E.C. provided resources and supervised experiments. C.D.C.B. supervised the entire study. A.V.P. and C.D.C.B. analyzed all results and wrote the manuscript. All authors have reviewed and approved the final manuscript.

